# Clustered and diffuse sensory corticostriatal wirings define complementary interneuron circuits

**DOI:** 10.64898/2026.10.04.756265

**Authors:** Juliette Contadini, Elodie Fino, Ingrid Bureau

## Abstract

In the striatum, interneurons tightly control the activity of output neurons, the striatal projection neurons (SPN). Among these striatal interneurons, parvalbumin-expressing (PV) and somatostatin-expressing (SOM) cells exhibit markedly distinct electrophysiological and anatomical properties, suggesting differential integration into striatal circuits and distinct forms of feedforward inhibition onto SPN. However, it remains unclear whether differences already emerge at the level of corticostriatal connectivity and if they shape specific interactions with cortical inputs to SPN. Here, we investigated the spatial organization of the projections from the primary and secondary somatosensory areas (S1 and S2), onto striatal PV and SOM interneurons and SPN. With a combination of anatomical tracing and functional circuit mapping, we characterized key features of corticostriatal connectivity, including input number, origin, spacing, overlap and strength. We found that input spacing and strength, at both the individual cell and population levels, were the principal features distinguishing PV and SOM interneurons. In contrast, connectivity differed primarily in input number and laminar organization across S1 and S2. Our findings suggest two potential modes of interaction between interneurons and SPN connectivity: one in which PV interneurons receive S1 inputs structured in large clusters, that densely overlap with those of neighboring SPN, providing an anatomical substrate for potential neuronal assemblies. And another in which SOM interneurons also receive abundant cortical inputs, but thinly disseminated across S1 and S2 and that overlap little with those of SPN, potentially shaping these cell integration through input cross-interaction. These cell-type principles of corticostriatal wiring may support distinct forms of striatal computation.

## INTRODUCTION

Striatal function critically depends on a small but diverse population of interneurons that exert powerful control over the striatal projection neurons (SPN). They have been implicated in the genesis of striatal activity underlying motor learning, automatized behaviors, and goal-directed action sequences (O’Hare et al. 2016; Martiros et al. 2018; Lee and Reeve 2017; Rotariu et al. 2025; Owen et al. 2018). Among striatal interneurons, the parvalbumin-expressing and somatostatin-expressing interneurons (termed PVi and SOMi thereafter) have rather distinct electrophysiological and anatomical features, leading to different feedforward inhibition onto SPN. More precisely, PVi display fast-spiking properties (Kawaguchi 1993; Koós and Tepper 1999; J. M. Tepper and Plenz 2006) and exert striatal region-specific modulation of SPN activity (Fino et al. 2018; Monteiro et al. 2018).

Anatomically, they are particularly numerous in the dorsolateral striatum (DLS) that integrates sensorimotor inputs (Gerfen 1985; Fino et al. 2018; Monteiro et al. 2018) and they present both dense and compact axon and dendrites, harboring more cortical synapses than other striatal neurons, (Kawaguchi 1993; Koós and Tepper 1999; Tepper and Bolam 2004). PVi-mediated feedforward inhibition of SPN is strong, fast and tightly controlling the timing and amplitude of their firing upon cortical inputs (Koós and Tepper 1999; Tepper et al. 2018). On the other hand, SOMi are electrically compact and more excitable than PVi, identifying them as potent modulators of striatal activity (Kawaguchi 1993; Tepper et al. 2010; Rotariu et al. 2025). While being more homogeneously distributed than PVi, region-specific excitability results in a weaker inhibition weight of SOMi on DLS SPN (Fino et al. 2018). Both their dendritic and axonal arbors are sparse and span a larger striatal territory, extending approximately twice as far as those of PVi (DiFiglia and Aronin 1982; Kawaguchi 1993; Kubota and Kawaguchi 2000; Tepper et al. 2010). This suggests that SOMi form different networks compared to PVi, perhaps reaching out to more distant pre- and postsynaptic cells. Overall, these distinct properties suggest that PV and SOM interneurons may be differently integrated into striatal networks and understanding the logic of their wiring could be central to assess their respective role in controlling striatal output.

We hypothesize that wiring differences between PV and SOM interneurons may already emerge at the level of projections connecting the sensorimotor cortical areas to the DLS. While cortical stimulations have provided initial evidence for such differences in projection strength, with PVi receiving stronger inputs than SOMi (Johansson and Silberberg 2020; Assous et al. 2018; Sharott et al. 2012), key features of connectivity patterns, such as the input cell laminar distribution and dispersion across cortical columns of the sensorimotor areas remain uncharacterized. Importantly, it is also unknown whether projections to interneurons and SPN are similarly organized and eventually shared. Yet, their degree of overlap is an essential issue for understanding the function of feedforward inhibition. A first characterization of SPN connectivity pattern in the primary somatosensory cortex (S1) showed that, although projections from multiple cortical columns converged onto the same region of DLS, individual SPN received largely distinct sets of cortical inputs, even when closely positioned in the DLS (Amroune et al. 2025). One may hypothesize that dense cortical inputs to interneurons, intersecting with those of many SPN, would enable scaled and temporally precise feedforward inhibition, thereby promoting the formation of functionally coherent assemblies. Conversely, non-overlapping connectivities could support other striatal computations, based on the interactions between distinct cortical inputs to interneurons and SPN. Finally, how generalizable connectivity patterns are across cortical areas is also a determinant question (Wall et al. 2013; Assous and Tepper 2019; Johansson and Silberberg 2020). Differences in selectivity and perceptual representations precision between S1 and the secondary somatosensory cortex (S2) (Carvell and Simons 1986; Kwegyir-Afful and Keller 2004; Hubatz et al. 2020; El-Boustani et al. 2020) may shape specific corticostriatal projections and connectivity pattern interactions in the DLS.

To test these hypotheses, we investigated the spatial organization of neurons in S1 and S2 projecting onto DLS PVi, SOMi and SPN using a combination of anatomical rabies-based retrograde tracing and functional mapping approaches. We found a profound difference in the spatial organization of both anatomical and functional projections targeting PV and SOM interneurons, particularly those originating from S1. They suggest two potential modes of interactions with SPN: One, in which PVi inputs, organized in clusters in S1, overlap with those received by a large population of SPN and the second in which sensory dispersed inputs to SOMi, but also sparse S2 inputs to PVi, may shape SPN integration by conveying inputs of different spectrum.

## RESULTS

### Distinct large scale spatial organizations of PVi and SOMi presynaptic cells

We first traced the presynaptic cells of PV and SOM interneurons in the somatosensory cortex using monosynaptic modified rabies virus. In SOM-cre and PV-cre mice, sequential unilateral injections of floxed TVA helper virus and glycoprotein (GFP fused N2cG) and rabies virus EnvA-CVS-N2c-ΔG (mCherry fused) were performed in the DLS (Fig. 1A-C). Systematic serial brain sectioning and 3D reconstruction allowed us to localize starter cells in the DLS and presynaptic cells in the somatosensory cortex. PV and SOM starter cells were similarly distributed within the DLS (Fig. 1D; n = 4 and n = 5 brains respectively). For both populations, repartition of presynaptic cells showed a strong bias toward the ipsilateral somatosensory cortex (PVi, 88 ± 1 %; SOMi, 93 ± 1 %). The number of presynaptic cells located ipsilaterally was tightly correlated with the number of PV starter cells (R^2^ > 0.99), corresponding to an estimated 24 presynaptic cells per PV starter cell in the ipsilateral somatosensory cortex (Fig. 1E). In contrast, no clear relationship was observed for SOM starter cells (R^2^ = 0.12), with the presynaptic-to-starter cell ratio ranging from 24:1 to 110:1 across brains. Nevertheless, presynaptic cell count did not scale differently with starter cell count between PVi and SOMi (interaction, *p* = 0.30).

**Figure 1.**
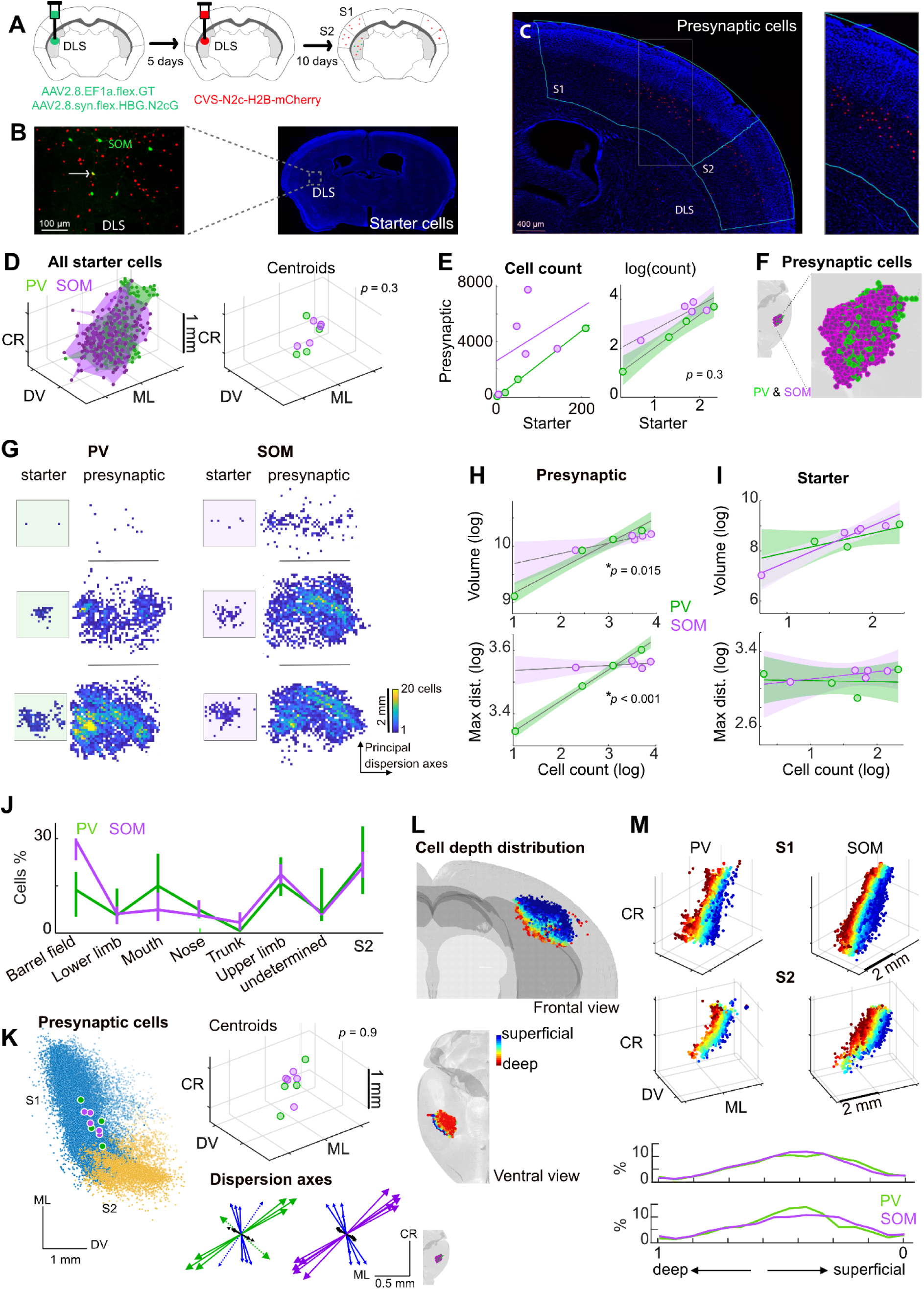
Distinct rules governing the spatial organization of PVi and SOMi presynaptic cells revealed with anatomical tracing. **A**: Experimental workflow showing sequential unilateral viral injections for cell type-specific monosynaptic modified-RV tracing of cells projecting to DLS. **B**: Left, magnification of the injection site in DLS. In green, GFP in the infected SOM cells, in red, mCherry in the striatal presynaptic cells and in yellow the co-infected SOM cells, i.e. the starter cells (arrow). Scale bar, 100 μm. Right, coronal section of a SOM-cre mouse, with DLS SOM cells infection and retrograde labelling of presynaptic cells. In blue, DAPI staining. Scale bar, 500 μm. **C**: Representative coronal section including S1 and S2 (cyan solid lines) at two magnifications. In red the cortical presynaptic cells, in blue the DAPI staining. Scale bar, 400 μm. **D**: The ensemble of starter PV (green) and SOM (purple) interneurons and each brain centroid. Green, PV, n = 4; Purple, SOM, n = 5. **E**: Number of presynaptic cells as a function of PV and SOM starter cells (each point represents one brain). Raw data (left) and log_10_ (right). Fits and 95 % confidence intervals are shown. **F**: The ensemble of presynaptic cells of striatal PV and SOM interneurons at two magnifications. **G**: Example two-dimensional histograms of starter and presynaptic cells along the two principal dispersion axes of cell distribution. The color codes for the number of cells in every 100 μm bin. Each row corresponds to a brain with either PV starter or SOM starter cells (left and right, respectively). Brains at the top and bottom are with the fewest and most starter cells, respectively. **H**: Relationship between cell count and population volume (top, log_10_(*μ*m^3^)) and mean maximal intercell distance (bottom, log_10_(*μ*m)) for the PVi and SOMi presynaptic populations. **I**: The same as in H for starter cells. One PV brain has 2 starters (top example in G), hence the 3 green symbols in the volume plot and 4 in the mean maximal intercell distance plot. J: Proportion of PVi and SOMi presynaptic cells in the different territories of the somatosensory cortex (median and 25-75th precentiles). **K**: Top, the centroids of the PVi (green) and SOMi (purple) presynaptic cells overlaid on all presynaptic cells, viewed in two dimensions. Cells in S1, blue, and in S2, yellow. Bottom, the three principal axes of cell dispersion (1st, green-PV or purple-SOM; 2nd, blue; 3rd, black), viewed from dorsal. The dotted-line arrows represent the brain with the fewest starter PVi (2) and presynaptic cells in the somatosensory cortex (11). **L**: All the presynaptic cells color-coded as a function of their positions along the third principal axis of cell dispersion, used as a proxy for cortical depth. Cells are shown in frontal and ventral views of the brain. **M**: Top, the same as in L, for PVi and SOMi presynaptic cells in S1 and S2 separately. Bottom, distribution of presynaptic cells along normalized cortical depth, with cortical depth ranging from 0 (superficial) to 1 (deep).

We next asked whether the organizations of PVi and SOMi presynaptic cells differed (Fig. 1F,G). To test whether they were distributed over territories of different sizes, we compared the relationships between presynaptic cell count and volume, and between presynaptic cell count and mean maximal intercell distance (Methods). Both relationships differed significantly between PVi and SOMi, with a larger difference observed for mean maximal intercell distance (Fig. 1H; interaction, *p* = 0.015 and *p* = 0.00015). This indicated stronger scaling of the spatial distribution of PVi presynaptic cells with cell number. In fact, mean maximal intercell distance of SOMi presynaptic cells showed little scaling with cell number, as illustrated in the examples of Fig. 1G. These differences could either reflect intrinsic properties of the presynaptic populations or arise from differences in starter cell organization. Against the latter possibility, neither relationship differed between the PV and SOM starter populations (Fig. 1I). These observations argue for genuine differences in the spatial distribution of presynaptic cells targeting PVi and SOMi.

We tested the possibility that these differences were due to distinct distributions of PVi and SOMi presynaptic cells across the somatosensory cortex. First, PVi and SOMi presynaptic cells were in similar proportions in the somatosensory cortex (16 ± 3% and 22 ± 1% respectively; *p* = 0.29). Both showed a similar bias toward S1 relative to S2 (77 % vs. 23 % and 78 % vs. 22 %; Fig. 1J). The centroids of PVi and SOMi connectivity were similarly localized within S1 (Fig. 1K, *p* = 0.88), and their principal axes of presynaptic dispersion were aligned (Fig. 1K, the arrows). Finally, PVi and SOMi presynaptic cells were both evenly distributed across S1 functional subdivisions (p = 0.89, Two-way Anova, Fig. 1J). Next, we compared the layer distributions of PVi and SOMi presynaptic cells (Fig. 1L; Methods). Likewise, no significant differences were found between the two interneurons populations in either S1 or S2 (Fig. 1M). Thus, PVi and SOMi connectivity did not differ in the global localization of presynaptic cells within the somatosensory cortex.

This suggested that the distinct rules organizing PVi and SOMi connectivity might also manifest at a finer spatial scale, within cortical subdivisions. However, whether these rules operated at the level of individual interneurons or instead organized connectivity at the population level remained unclear. Indeed, the weaker scaling of the SOMi presynaptic population volume could reflect either individual interneurons sampling inputs in broader cortical territories than PVi neurons, or SOMi neurons sampling inputs in more overlapping cortical territories. We therefore tested these two possibilities by examining the spatial organization of connectivity at the level of individual interneurons using LSPS mapping and single-cell recordings.

### Clustered PVi connectivity vs. diffuse SOMi and SPN connectivity in the barrel field

From PV-cre::ai14 and SOM-cre::ai14 mice, we prepared slices preserving the corticostriatal projections originating from one row of cortical columns in the barrel field (Amroune et al, 2025), then uncaged glutamate with laser-scanning photostimulation (LSPS; 2.1 × 1.1 mm, 75 μm spacing; Methods) at different sites to stimulate cortical cell bodies and recorded the evoked synaptic responses in mCherry-labeled striatal PV or SOM interneurons (Fig. 2A; Fig. S1A). In a subset of experiments, recordings of nearby SPN were performed to allow direct paired comparisons. Connectivity maps identified the sites where LSPS evoked EPSCs (termed connecting sites), thereby revealing the locations of presynaptic excitatory neurons (Fig. 2B,C). PVi had twice the number of these connecting sites compared to SOMi and same slice SPN (Fig. S1B,C). Their inputs were also individually stronger compared to SOMi (Fig. S1F,G). To test the hypothesis of different input spatial organizations, we measured for each interneuron recording the width of the entire cortical stretch containing its connecting sites, termed input field hereafter, the number of cortical columns and layers in which these sites were distributed. Irrespective of these anatomical landmarks, we also measured the number and size of clusters formed by the connecting sites (Fig. 2D; Methods). Multivariate analysis revealed significant differences across cell types, with PVi connectivity maps segregating from those of SOMi and SPN (permanova, pseudo-*F* = 17.2, *p* = 0.0005; Fig. 2E; Methods). The variables contributing most strongly to this separation were the number of cortical columns, the number of layers, and the size of the largest cluster, all of which were greater for PVi (Fig. 2F). Notably, the greater number of connecting sites in PVi connectivity maps was not associated with broader input fields or a larger number of clusters than in SOMi maps (Fig. 2D). Instead, PVi connecting sites were organized into larger clusters. Consistent with this, the largest PVi clusters spanned nearly three barrel-columns along the cortical horizontal axis (413 ± 38 μm), whereas the largest SOMi and SPNi clusters extended only about half this distance (Fig. 2G; Kruskal–Wallis, *p* < 0.0001; PV-SOM, *p* = 0.022; PV-SPN, *p* < 0.001, Dunn-Šidák).

**Figure 2:**
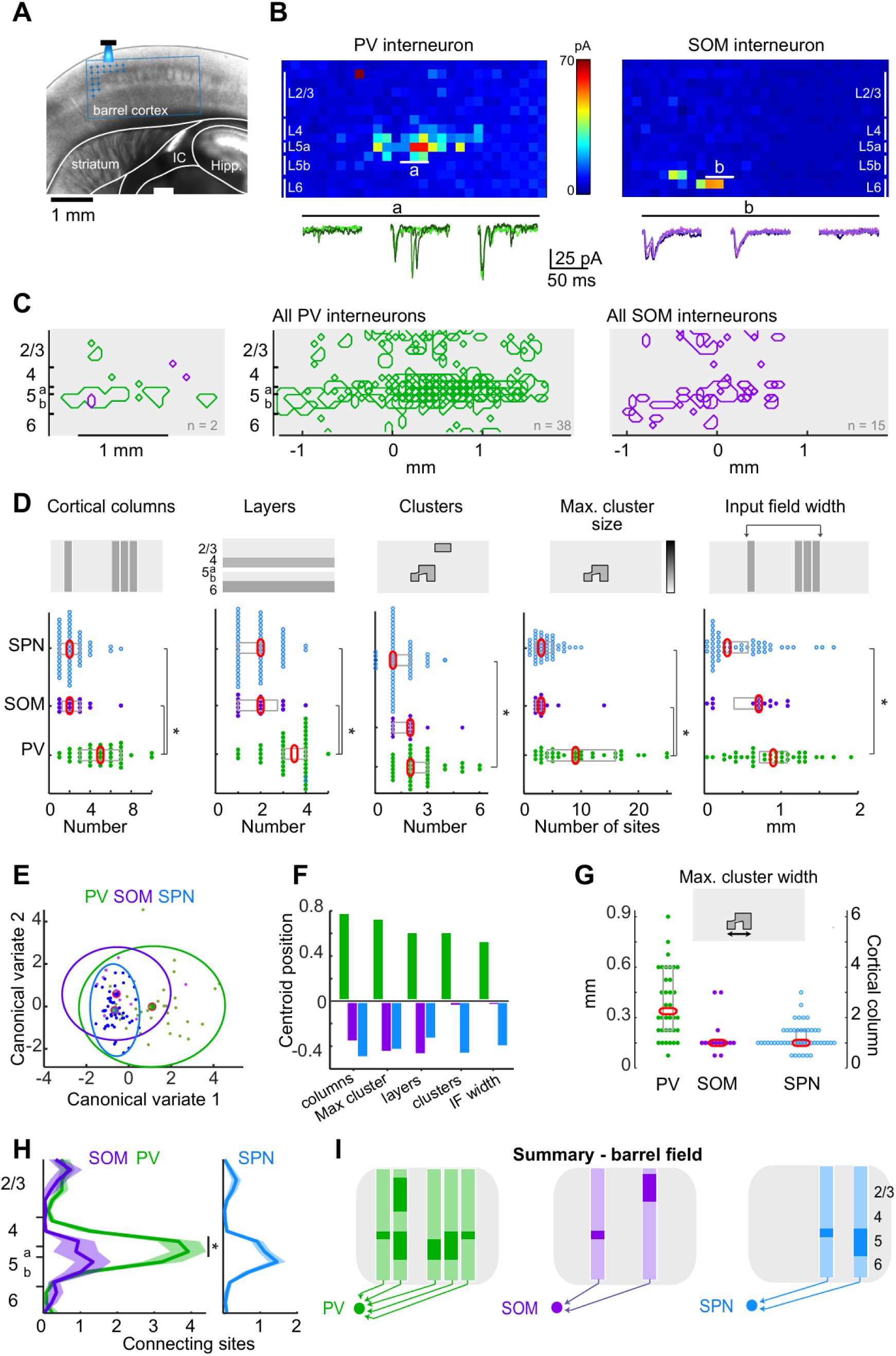
Clustered PVi connectivity vs. diffuse SOMi and SPN connectivity in the barrel field revealed by single cell recordings and LSPS mapping. **A**, Montage of a DIC image of a barrel cortex corticostriatal slice with superimposed schematic representation of the experiment. Striatal cells (PVi, SOMi or SPN) were recorded in the DLS while LSPS with glutamate uncaging was performed in the barrel cortex (blue grid). IC, internal capsule; hipp., hippocampus. **B,** Top, examples of synaptic input maps for individual PV and SOM interneurons. Each pixel of color indicates the peak amplitude of EPSCs evoked by LSPS in the barrel field. The cortical layers are indicated by solid white vertical lines. Bottom, EPSC traces evoked at the sites indicated by letters in the maps above. Four repetitions are superimposed. **C,** Left Example connectivity maps (polygons representing connecting sites) of one PV (green) and one SOM (purple) interneuron. Right, overlay of sites in the barrel cortex where stimulations evoked EPSC in PV (n = 38) and SOM (n = 15) interneurons. **D:** Number of cortical columns, layers, clusters, size of largest cluster and input field width in connectivity maps of PVi, SOMi and SPN (n = 50, of which 35 were in Amroune et al., 2025). *, Columns: PV-SOM: p = 0.0013, PV-SPN: p < 0.001; Layers: PV-SOM: p = 0.0013, PV-SPN: p < 0.001; Clusters: PV-SPN: p < 0.001; Max cluster size: PV-SOM: p < 0.001, PV-SPN: p < 0.001; input field width: PV-SPN: p < 0.001; Kruskal-Wallis, FDR correction. **E**, Canonical variate analysis of connectivity parameters across neuronal types, the first two canonical variates. Ellipses show 95 % confidence regions for each group, and large red symbols indicate group centroids. (permanova, pseudo-F = 17.2, p < 0.0005). **F**, Relative contribution of each parameter to PV-SOM-SPN centroids separation. **G**, Width of the largest cluster. **H**, Number of connecting sites across cortical layers. *, p = 0.0012 and < 0.001; permutation-based test with FDR correction. I, Schematised median connectivity patterns in the barrel field.

The number of innervating cortical layers being one distinctive characteristic of PVi, we further studied the connectivity maps along their vertical axis. Like previously found for SPN (Amroune et al., 2025), L5 dominated the inputs to both PV and SOM interneurons. However, connecting sites were ∼4 fold denser in L5 in PVi maps (Fig. 2H; *p* < 0.05; permutation-test). Thus, inputs to individual PVi appeared particularly aggregated in L5, and those to SOMi more broadly distributed throughout L2/3, 5 and 6 (Summary in Fig. 2I).

These results showed that PVi had individually more inputs present in the slice than SOMi, and that these formed compact clusters. To compare the cortical input spaces between individual interneurons, we analyzed the connectivity maps that were obtained for several PVi or SOMi in the same slice (see Methods). This showed that a large majority of neighboring PVi had their connecting sites located in one or several common clusters in the barrel field (78 %; n = 35 pairs) (Fig. 3A-B). This rate decreased slowly with interneuron spacing, as sharing of connecting sites was possible for up to an estimated PVi separation of 1.5 mm in the striatum. On average, each PVi pair had clusters of shared connectivity constituted by 2.3 ± 0.3 connecting sites, but these could occasionally reach 12 sites (Fig. 3C). In contrast, consistent with their sparser innervation, only 25 % of SOMi pairs had clusters of shared connectivity (2 of 8 pairs), and these were both less numerous and smaller, generally consisting of a single site (1.2 ± 0.1). Thus, functional inputs to SOMi were not aggregated into hotspots of connectivity. These observations suggest that, within the reduced dimensionality of a single row of cortical columns, SOMi predominantly sampled distinct territories whereas PVi tended to sample overlapping territories.

**Figure 3.**
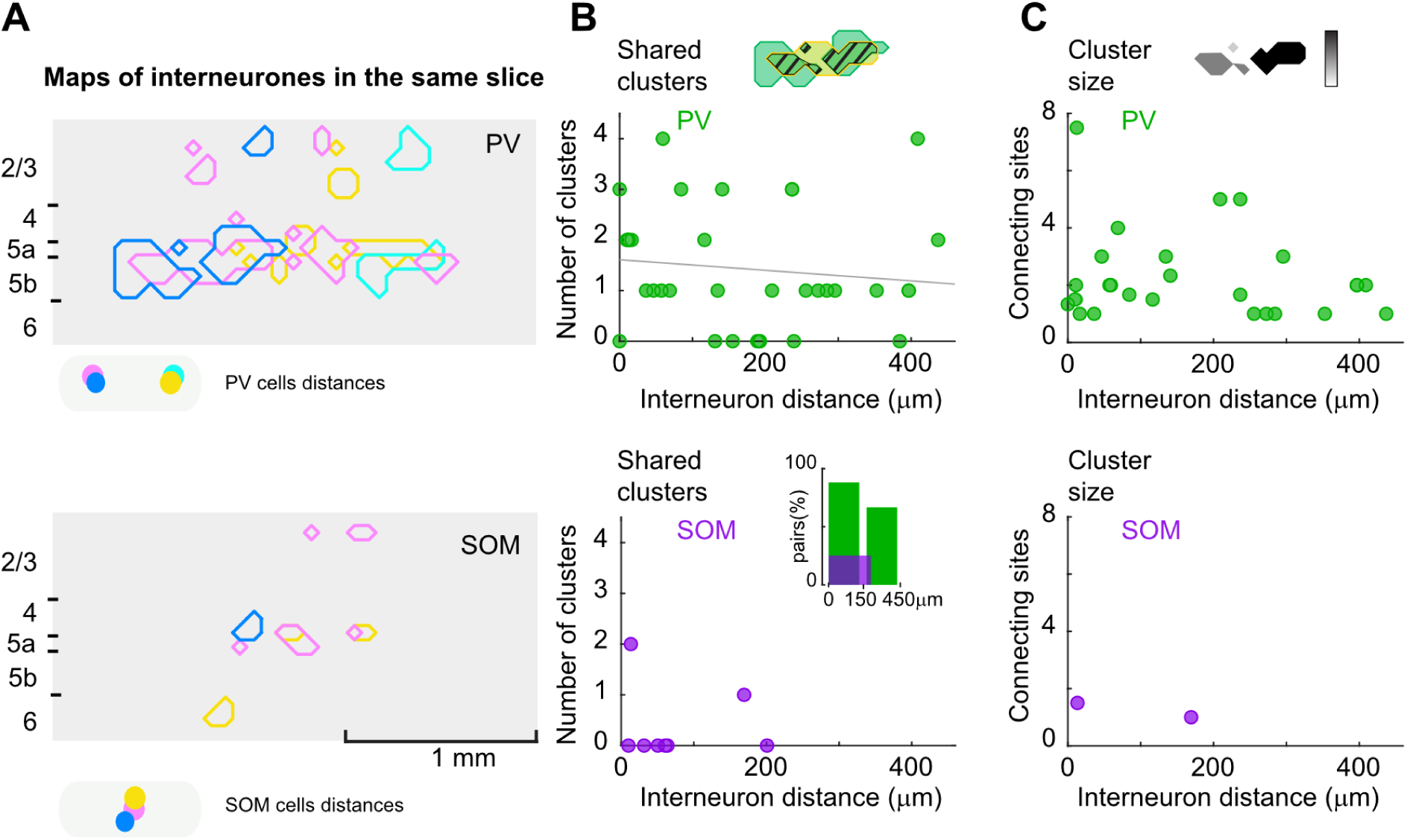
Co-clustering of PVi and spatial dispersion of SOMi connecting sites. **A**, Connectivity maps of interneurons, PV (top) or SOM (bottom) interneurons, recorded in the same slice, in the barrel field. At the bottom, filled symbols show the distances between interneurons in the striatum. Neurons are color coded. **B**, Number of input clusters shared among interneurons in each pair for PVi (top, n = 35 pairs) and SOMi (bottom, n = 8) as a function of cell distance. Bars in the inset show the proportions of PV or SOM pairs sharing connecting sites. **C**, Number of connecting sites constituting the shared clusters for PV (top) and SOM (bottom) pairs as a function of cell distance.

### S2-specific functional connectivity patterns

Our results showed that the spatial organization of functional inputs from the barrel field differed significantly between PV and SOM interneurons. We next investigated whether this was generalizable to projections from S2. Single cell recordings and LSPS were repeated in a preparation containing both S2 and its corticostriatal projections (Fino et al., 2005) and the same organization parameters were evaluated as before (Fig. 4A-D). A first striking observation was the number of connecting sites present in the PVi connectivity maps which was ∼ 2 fold lower compared to the barrel field (7.7 ± 1.6, n = 15; Fig. 4C; Suppl. Fig. 2D). There was a tendency for PVi to still have more connecting sites than SOMi but the difference was not significant (SOMi: 3.6 ± 0.9, n = 9; p = 0.06, Kruskal-Wallis, Fig. S1D). In fact, large clusters were no longer a characteristic of PVi connectivity patterns, now 170 ± 23 µm for the largest within each map, or ∼1 cortical column (Fig. 4E). The consequence of this lower connectivity could also be measured in that connecting sites in S2 were never shared among PVi (n = 6 pairs). A second feature of S2 connectivity maps was the major decrease in L5 inputs. This was visible in the PVi inputs more homogeneously distributed along the vertical axis compared to the barrel field, suggesting that inputs were principally shed of layer 5 (Fig. 4F). Nevertheless, multivariate analysis also revealed in S2 significant differences between the spatial organizations of the PVi, SOMi and SPN inputs (*p* = 0.044, pseudo-F = 2.6), only this time SOMi showed the greatest separation (Fig. 4G). Strongly contributing to their separation was SOMi input distribution in less cortical columns and smaller input fields compared to PVi and SPN (Fig. 4D). These results suggest that, within globally sparse S2 projections, SOMi functional connectivity was distinctively low (summary in Fig. 4H).

**Figure 4:**
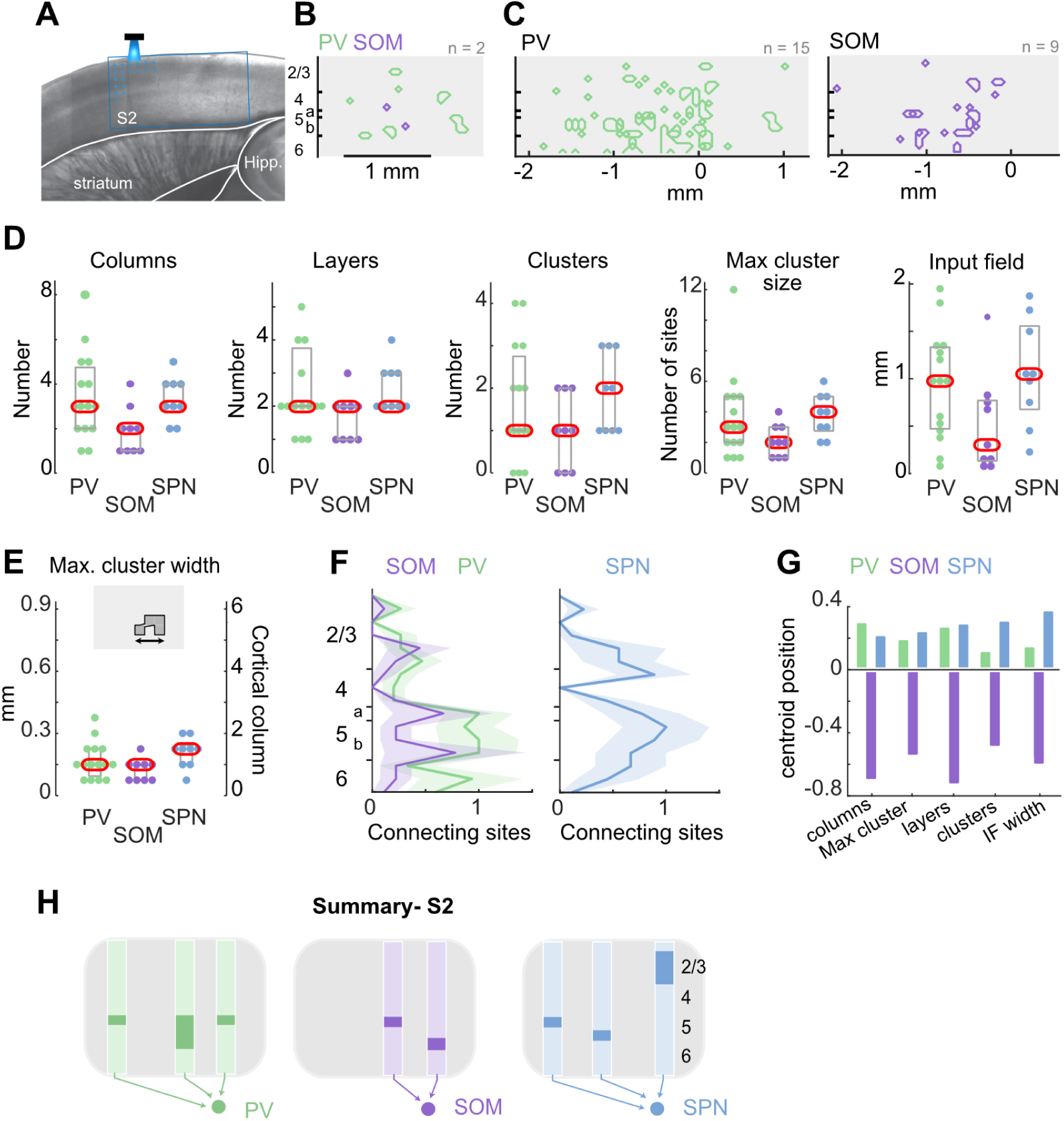
S2-specific distinction between interneurons functional connectivity patterns. **A**, Montage of a DIC image of a S2 corticostriatal slice with superimposed schematic representation of the experiment. Striatal cells (PV, SOM or SPN) were recorded in the DLS while LSPS with glutamate uncaging (blue grid) was performed in S2. **B**, Example connectivity maps (polygons) of one PV (green) and one SOM (purple) interneuron. **C**, overlay of sites in S2 where stimulations evoked EPSC in PV (n=15) and SOM (n=9) interneurons. **D**, from left to right, number of cortical columns, layers, number and maximal cluster size, and input field width connected to PVi (n=15), SOMi (n=9) and SPN (n=9) cells. **E**, Width of the largest cluster. **F**, Number of connecting sites across cortical layers. **G**, Relative contribution of each connectivity parameter to PV-SOM-SPN centroids separation in canonical variate analysis across neuronal types (permanova, p = 0.044, pseudo-F = 2.6). **H**, Schematised median connectivity patterns in S2.

### Different degrees of overlap between interneurons and SPN inputs

Finally, to evaluate the potential consequences of specific PVi and SOMi innervation patterns on the control of striatal somatosensory integration, we compared the connectivity maps between nearby SPN and interneurons recorded in the same slice (Fig. 5A). We measured both the vertical alignment (Fig. 5B and D) and the 2D overlap (Fig. 5C and E) of SPN and interneuron connecting sites. Whereas vertical alignment reflects whether inputs arise from the same whisker column, 2D overlaps incorporate the layers which are associated with distinct sensory integration pathways (El-Boustani et al. 2020; Bureau et al. 2006). We found that nearly all SPN and PVi positioned within 150 µm, and still 67 % of pairs within 300 µm separation, had connecting sites vertically aligned (Fig. 5B; n = 19 pairs). Overall, as much as 45 ± 7 % of SPN connecting sites were in the same cortical columns as those of a PVi. The chance of these sites being in the same layer too was also high as 2D overlap was observed in 90-60 % of the pairs and still for 24 ± 5 % of the SPN overall connectivity (Fig. 5C). Interestingly, despite SOMi connecting sites being more scattered, their vertical alignment with those of SPN was nevertheless high, observed in 4 out of 5 SOM-SPN pairs (<175 µm distance; Fig. 5B). However, 2D overlap was less frequent (2 out of 5 pairs) and included only 6.7 ± 4.4 % of the SPN connecting sites. This suggested larger input divergence in the laminar axis between SOMi and SPN compared to the PV-SPN pairs (Fig. 5C; summary in Fig. 5F).

**Figure 5:**
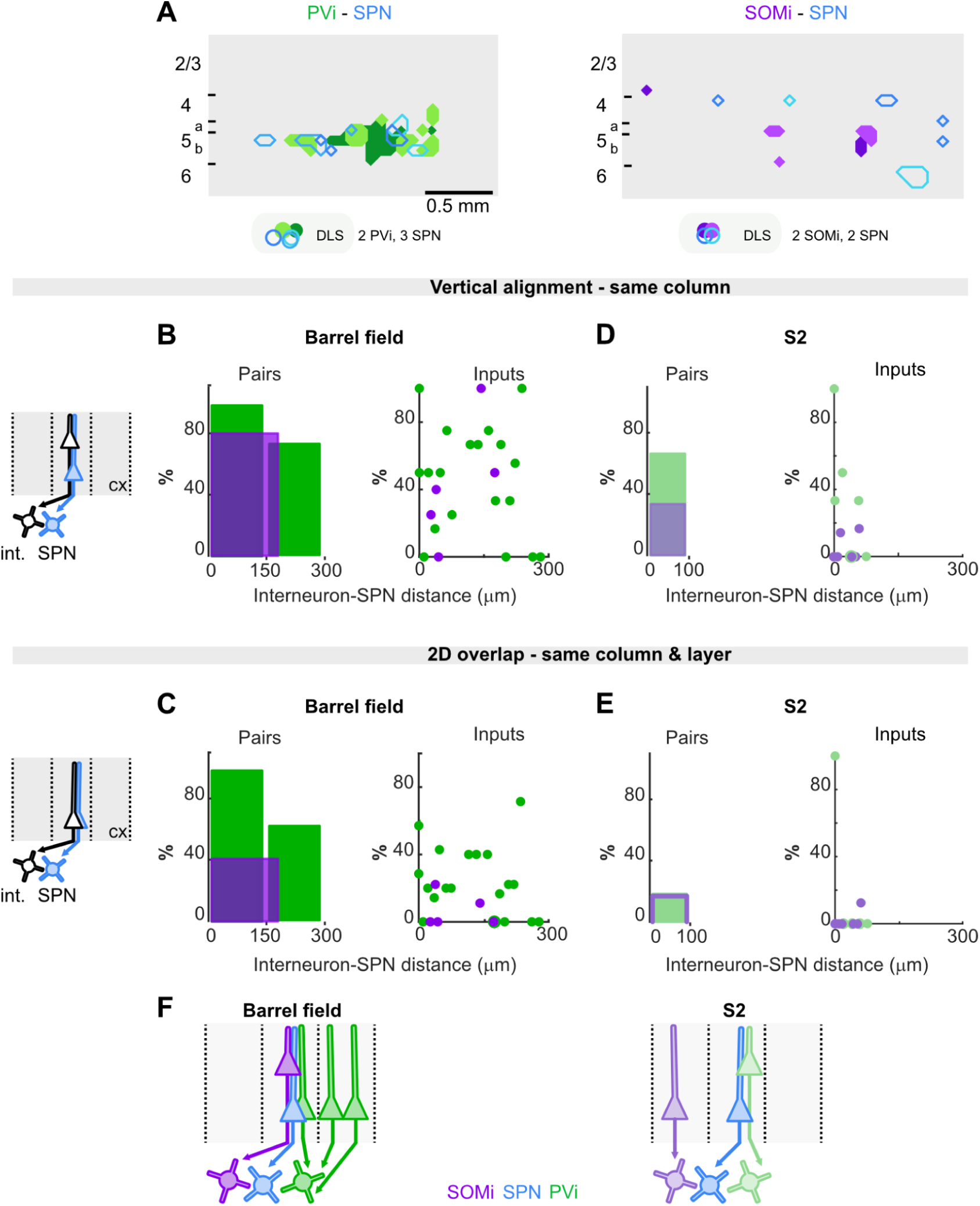
The projections to PV and SOM interneurons overlap to different degrees with projections to SPN. **A**, Top, connectivity maps of interneurons and SPN recorded in the same slice, in the barrel field or in S2. Bottom, cell distances in the striatum. PV, green shades; SOM, purple shades; SPN, blue shades. **B**, Left, In the barrel field, percentage of PV-SPN pairs (n = 19) and SOM-SPN pairs (n = 5) with overlapping columns for increasing distances between cells. Right, percentage of SPN columns overlapping with those of PV or SOM interneurons. **C**, same as in B for the SPN connecting sites overlapping with those of interneurons. Right, % of pairs. Left, % of SPN connecting sites. **D,E** same as in B,C for S2. PV-SPN pairs, n = 5; SOM-SPN pairs n= 6. **F**, Schematized positions of the interneurons inputs relative to one given SPN input in the barrel field (left) and in S2 (right).

A different picture emerged from the connectivity maps in S2 where overlap could occur for similar long distances separating cells, but in a sparse manner for both PV and SOM interneurons (Fig. 5D). The chance that neighboring PVi and SPN had connecting sites vertically aligned in S2 was still relatively high, observed in 4 out of 6 pairs (Fig. 5D), but unlike in the barrel field they were rarely in the same layer (in 1 pair; Fig. 5E). Finally few SOM-SPN pairs had connecting sites vertically aligned or in common in S2 (1 out of 6 pairs; Fig. 5A,B,D; summary in Fig. 5F).

## DISCUSSION

We investigated the organization of somatosensory corticostriatal projections targeting the PV and SOM interneurons and the SPN in the DLS and found that innervation patterns varied both as a function of cell-type and somatosensory cortical area. S1 projections to PV and SOM interneurons were both prominent, but their spatial organization differed significantly: presynaptic cells targeting SOMi were dispersed, whereas those targeting PVi formed clustered distributions. The projections from the barrel field targeting the PVi overlapped with those targeting the nearby SPN and other PVi. In contrast, the projections to SOMi rarely overlapped with those to SPN and other SOMi. These distinct connectivity patterns suggest the existence of orthogonal striatal computations through cell- and area-specific interactions between the interneurons and SPN.

### Sensory cortical connectivity positions PV interneurons as organizers of striatal activity in the DLS

The connectivity pattern of PVi may represent an important determinant of their rapid control over SPN populations. PVi exhibited a stereotyped and clustered organization of their presynaptic inputs within S1, supported by several independent observations. At the population level, anatomical tracing revealed a strong coupling both in size, between PV starter and presynaptic ensembles, and in space, between presynaptic cell number and occupied space in the somatosensory cortex. Investigated with LSPS mapping in a reduced dimensional space, connectivity patterns consisted in discrete but relatively large clusters of functional inputs, predominantly in L5. LSPS also revealed substantial overlap in cortical inputs among neighboring PVi that was not suspected from the strong linear relationship between starter and presynaptic cell counts in the tracing data. It suggests that neighboring PVi sample inputs in the same precise cortical territories, not from the same cortical cells. This clustered organization contrasts with the more dispersed and heterogeneous connectivity observed for SOMi and SPN populations.

Another prominent feature of PVi connectivity in the barrel field was its substantial overlap with SPN inputs. Nearly all neighboring PVi and SPN shared cortical columns, and up to one quarter of the connecting sites in a SPN map overlapped with those of a single PVi, indicating that these inputs originated from the same precise cortical locations. This extensive overlap is favored by PVi input clusters spanning multiple cortical columns and layers. Such an organization is well suited to the co-activation of individual PVi with multiple SPN despite the heterogeneity of SPN cortical inputs (Amroune et al. 2025). Consistent with this interpretation, comparisons across multiple LSPS recordings showed that the input sites of individual PVi overlapped with those of several SPN, and conversely that individual SPN shared input sites with multiple neighboring PVi. Together, these findings suggest that much of the cortical input received by a SPN is paralleled by the combined cortical inputs to the surrounding PV population. This organization appears specific to the PV-SPN relationship, as neighboring SPN share relatively few cortical inputs with one another (Amroune et al. 2025), suggesting distinct functional roles for inhibition between SPN and feedforward inhibition mediated by PVi. Combined with the strong efficacy of PVi→SPN synapses (Kawaguchi 1993; Tepper et al. 2010; Klaus et al. 2017) and the reliable, time-locked recruitment of PVi by cortical activity (Mallet et al. 2005; Pidoux et al. 2011;Fino et al. 2018; Druart et al. 2025), this precise alignment of cortical inputs provides an anatomical substrate for feedforward inhibition of SPN with millisecond precision. At the population level, the simultaneous convergence and divergence of cortical input onto PVi is likely to sharpen the temporal control of SPN ensembles and thereby promote the emergence of coordinated striatal assemblies (Costa et al. 2006; Moyer et al. 2014; Barbera et al. 2016; Burke et al. 2017) whose activity would be proportionally more shaped by the cortical inputs to PVi.

However, LSPS mapping unveiled areal specificities which refined the function of PVi in the DLS: PVi inputs in S2 were sparse, less focused on L5, and different from those of neighboring SPN and other PVi. This sparse connectivity of PVi together with the different laminar organization and the relatively wider extent of SPN connectivity (Fig. 4D,F) could reflect the distinct properties of S2 which as a second order area integrates multiple perceptual representations (Taub et al. 2024; Timmermann et al. 2001; Hubatz et al. 2020) and whose outputs may already combine diverse information. We previously showed that the strength of inhibition onto SPN was heterogeneous across striatal regions (Fino et al. 2018). The present study suggests an additional level of refinement to feedforward inhibition whereby cortical connectivity of PVi is scaled in accordance to the intrinsic characteristics of the cortical areas. This also implies that other forms of inhibition may have greater weight over the SPN integration of S2 inputs.

### Widespread sensory cortical connectivity positions SOM interneurons as broad spectrum integrators in the DLS

Retrograde tracing revealed a large number of presynaptic cells in the somatosensory cortex, identifying SOMi as a major target of corticostriatal projections. At first glance, this appears inconsistent with the weak inputs reported in previous electrophysiological studies (Johansson and Silberberg, 2020; Gittis et al., 2010) and with the sparse functional connectivity to SOMi revealed by LSPS here. However, these findings can be reconciled by two complementary factors. First, the weaker synaptic strength of corticostriatal connections onto SOMi, together with their occasionally distal dendritic location, may reduce their detection (Fig. S2G; Vuillet et al. 1992; DiFiglia and Aronin 1982; Kawaguchi 1993; Kubota and Kawaguchi 2000). Second, and more novel, the spatial organization of SOMi anatomical connectivity may contribute to the apparent discrepancy: if each SOMi samples inputs distributed across a cortical territory substantially larger than the one contained in *ex vivo* preparations, the mapping would only capture a smaller fraction of their presynaptic network relative to PVi. This interpretation is consistent with the weak relationship between SOM starter and presynaptic cell numbers which suggests that the number of cortical inputs received by SOMi may not be constrained independently within each cortical territory, but rather across a broader cortical scale.

Our results identify organizational principles for SOMi inputs that contrast sharply with those of PVi and suggest a distinct role in striatal computations. At the population level, SOMi presynaptic cells were distributed over larger cortical volumes compared to PVi. Examined at a lower scale and functionally with LSPS, individual SOMi connectivity patterns consisted in small clusters, approximately the width of a single cortical column, yet distributed across comparatively broad input fields relative to PVi. In addition, input sites were rarely shared between neighboring SOMi, suggesting that the sparse functional innervation of SOMi was not particularly biased toward specific anatomical connectivity sites in the barrel cortex. Instead, these results reveal an organization by which neighboring SOMi share large cortical input space wherein each interneuron strongest inputs are thinly disseminated. This pattern implies a mode of integration in which SOMi sample distributed cortical activity rather than being dominated by a specific input channel. It does also raise the intriguing possibility that such a large pool of ‘latent’ cortical inputs may undergo targeted synaptic plasticity, as observed in the dorsomedial striatum after learning a sensorimotor task (Rotariu et al. 2025).

SOMi are rarely contacted by PVi (Szydlowski et al. 2013; Gittis et al. 2010) suggesting that they operate a parallel control over SPN activity. With millimeter long axons (DiFiglia and Aronin 1982; Kawaguchi 1993; Kubota and Kawaguchi 2000), SOMi seem well positioned to functionally link together SPN distributed in different striatal regions. Our findings are aligned with this idea of large-scale modulator and go further showing that SOMi share almost none of their functional inputs with neighboring SPN: these were from a different layer in the barrel field and altogether from a different column in S2. These properties suggest a model in which SOMi modulate SPN activity through cross-interactions between distinct cortical inputs. Hence, DLS SOMi could shape the sensory context in which striatal computations unfold rather than enforcing coherent sensory tuning in SPN assemblies.

In conclusion, the distinct cortical connectivity patterns of PV and SOM interneurons, one clustered and stereotyped, the other diffuse and heterogeneous, may organize somatosensory integration in the DLS through two interwoven processes operating at complementary spatial scales. Together, these connectivity architectures could support the integration of both primary sensory inputs and higher-order sensory associations.

## MATERIEL ET METHODES

### Mice and ethics

PV-Cre and SOM-Cre lines were used for these experiments (mouse lines #008069 and #013044 from Jackson Laboratory), mated with ai14 mice (Jackson 007909). Both sexes were used on postnatal day 30 to 45 and housed in temperature-controlled rooms with standard 12 h light/dark cycles. Food and water were available *ad libitum*. All experiments were performed in accordance with EU institutional guidelines and French national institutional animal care guidelines (protocol #33185 and 27242).

### Pseudo rabies virus-based tracing

#### Stereotaxic injections

Animals were anesthetized with isoflurane (5% for induction and 2.5% for maintenance) and mounted on a stereotaxic device (Kopf Instruments; Tujunga, CA). After applying an external antiseptic and antifungal (Vetedine) and injection of local analgesic under the scalp, an incision was made along the midline of the skull. PV-cre and SOM-cre mice were injected with pseudo rabies virus to map inputs of either cell type. We previously confirmed the efficient and specific cre-dependent labeling with either GFP or mCherry in each interneuron using those cre-mouse lines (Fino et al. 2018; Rotariu et al. 2025). A small hole was drilled at the left dorsolateral (AP +0.8; ML +2.3; DV −2.5). 200nl of a mix of viruses (AAV2.8.EEF1a.flex.GTc and AAV2.8.syn.flex.H2BG.N2cG) were injected through a pulled glass pipette (pulled with a P-97 model Sutter Instrument Co. pipette puller) using a nanoinjector (World Precision Instruments, Germany) at a rate of 2 nl/sec. The injecting pipette was left in place for 2 to 3 minutes and then rapidly retracted to limit viral backflow. Mice were sutured and sterile saline was subcutaneously injected to minimize dehydration. After post-operative monitoring on a heating pad, mice were placed back to their home cage and monitored daily for 5 days. Five days later, 200 nl of modified rabies virus (RABV41-mCherry) were injected at the same coordinates in the same conditions. Mice were then monitored in their home cage during 10 days of viral expression. Different dilutions of the first mix of AAVs and RAB41 allowed to vary the number of starter cells.

#### Histology and microscopy acquisitions

Ten days after injection, mice were anesthetized with an overdose of zoletil/domitor (20 µl/10 g) and transcardially perfused with phosphate buffered saline (PBS) followed by 4 % paraformaldehyde (AntigenFix, Diapath) with a peristaltic perfusion pump. The brain was carefully extracted and post-fixed in 4 % PFA (48 hr). The brain was then transferred to PBS and kept at 4°C. Whole-brain coronal slices (50 µm) were collected with a vibratome (Leica) and simultaneously plated on super frost plus slides (Thermo Fisher Scientific, Waltham, MA). Slides were counterstained with DAPI (0.8 µg/ml) and cover slipped with mounting media containing (in mM): 115 Tris, 1870 Glycerol and 11.5 % Mowiol. When mounted, slices were acquired with a widefield microscope (DM6000, Leica) and a 10x objective. Acquisitions are done at three wavelengths: 358 nm (for DAPI staining), 395 nm (GFP, labeling of primary infected PV and SOM interneurons) and finally 550 nm (mCherry, labeling of presynaptic cells).

#### Brain slices alignment on Allen Brain Atlas

Acquired slices are then aligned to a reference coronal mouse brain atlas (Allen Brain Atlas, Allen Institute for Brain Science) using the Aligning Big Brains & Atlases plugging (EPFL,Chiaruttini et al. 2025) and its alignment protocol. Once slices are aligned, the reference atlas is transformed through manual and automatic procedures in order to fit optimally to the acquired slices.

#### Semi-automatized cell detections and quantifications

After atlas transformation, slices were exported to a Qupath project (Queen’s University Belfast, Northern Ireland) together with associated aligned atlas. Then starter cells and presynaptic cells were detected using a positive cell detection tool based on thresholded values of cells morphological features, and this for all channels, in a list of brain regions of interest. Starter cells were defined as detected co-infected cells with identical coordinates in both fluorescence channels and presynaptic cells were defined as cells detected in mCherry-channel.

#### Analysis of starter cells and presynaptic cells spatial organization

Mean maximal intercell distance is the maximal euclidean distance separating cells computed for each cell and averaged across the population. Principal component analysis (PCA) was performed on the xyz coordinates of presynaptic cells to study their principal axes of dispersion. To estimate the relative cortical depth of presynaptic cells, PCA was performed on the xyz coordinates of all PVi and SOMi presynaptic cells together to define a common anatomical reference frame. The third principal component (PC3), corresponding to the axis of lowest variance in the cell distribution and visually best aligned with the cortical depth axis (Fig. 1K,L), was used as an estimate of cortical depth. To account for local cortical curvature, the first two principal components (PC1 and PC2) were partitioned into 700 µm bins. Within each local PC1-PC2 bin, PC3 values were normalized between the 1st and 99th percentiles of the local distribution, thereby defining a relative cortical depth scale. This step was performed separately for S1 and S2. Each presynaptic cell was then assigned a relative depth value based on its normalized position along PC3. Cells located in sparsely populated bins were assigned depth values by nearest-neighbor interpolation from neighboring regions.

#### Brain slices preparation and electrophysiology

Mice were anesthetized with isoflurane (4 %) prior to cervical dislocation and decapitation. Corticostriatal slices of 350 μm thickness were cut in a chilled cutting solution containing (in mM): 110 choline chloride, 25 NaHCO_3_, 25 D-glucose, 11.6 sodium ascorbate, 7 MgCl_2_, 3.1 sodium pyruvate, 2.5 KCl, 1.25 NaH_2_PO_4_, and 0.5 CaCl_2_ (Sigma Aldrich). Barrel field-corticostriatal slices were cut with a 60° angle from the midline and a 10° angle in the dorso-ventral axis (Amroune et al., 2025). S2-corticostriatal slices obtained were horizontal (Fino et al., 2005). Slices were then placed in artificial cerebrospinal fluid (ACSF) containing (in mM): 127 NaCl, 25 NaHCO_3_, 25 D-glucose, 2.5 KCl, 1 MgCl_2_, 2 CaCl_2_, and 1.25 NaH_2_PO_4_, aerated with 95% O_2_ and 5% CO_2_. Slices were then transferred in an incubator at 34 °C for 30 min and then maintained at room temperature for 20 min prior to use. PV and SOM interneurons were visualized under infrared and fluorescent lights under a BX61WI microscope (Olympus) and patched with borosilicate electrodes (3-6 MΩ). Whole-cell recordings were performed in voltage-clamp using a Multiclamp 700A amplifier (Axon Instrument, Molecular Devices). The holding membrane potential was −70 mV for interneurons or −80 mV for SPN. The intracellular solution contained (in mM) 128 Cs-methylsulfate, 4 MgCl_2_, 10 HEPES, 1 EGTA, 4 Na_2_ATP, 0.4 Na_2_GTP, 10 Na-phosphocreatine, 3 ascorbic acid; pH 7.25; 290-300 mOsm. Non fluorescent cells were considered as SPN.

#### Laser scanning photostimulation (LSPS) with glutamate uncaging

LSPS was performed as previously described (Amroune et al, 2025). Slices were kept during recordings in circulating ACSF solution containing (in mM): 0.2 MNI-caged glutamate (Tocris), 0.005 CPP [()-3-(2-carboxypiperazin-4-yl) propyl-1-phosphonic acid], 4 CaCl_2_, and 4 MgCl_2_. Focal photolysis of caged glutamate was accomplished with a 2 ms 25 mW pulse of a pulsed UV (355 nm) laser (DPSS Lasers Inc.) through a 0.16 NA 4 × objective (Olympus). 464 positions spaced by 75 μm on a 29 × 16 grid (2.1 × 1.1 mm) were scanned over the cortex. The slice and the LSPS grid were oriented in such a way that the barrels were laid out horizontally. UV stimuli were applied every 700 ms and their successive positions on the LSPS grid were such as to maximize the time between stimulations of neighboring sites. Electrophysiological traces consisted of a 100 ms baseline prior to the 2ms laser pulse and 450 ms later a −5 mV 100 ms test pulse. 3 to 4 stimulations were performed at each site at ∼3 minutes intervals. Custom software for instrument control and acquisition (Suter et al. 2010) was written in Matlab (Mathworks). An image of the slice containing the stimulated barrel cortex and the connected DLS was acquired after each recording to report the position of all connecting sites and recorded cells in a common horizontal axis for every experiment.

#### Analysis of electrophysiologIcal recordings

Synaptic input maps of neurons were constructed by taking the peak amplitude of EPSC detected in a 50 ms time window starting at stimulation onset for each position in the LSPS grid. After a first round of automatic detection, traces were then visually inspected to detect small EPSC (overall range: 7 - 222 pA; mean 40 pA). To disambiguate evoked responses from spontaneous activity, EPSC occurring less than 2 to 3 times across repetitions (2-5) of maps were set to zero. Measures of genuine EPSC were then averaged across repetitions. Pattern analyses were written in Matlab (Bureau, 2025). We used the binary version of the LSPS map (‘connectivity map’) reporting the location of connecting and non-connecting sites (i.e., yielding an EPSC or none in the recorded cells). A cluster is constituted of 2 or more contiguous connecting sites. The number of cortical columns in the connectivity maps is derived based on the position of the connecting sites on the horizontal axis, considering the width of a barrel (150 μm in this preparation) and the spacing in the LSPS grid (75 μm). Hence, 1 cortical column is constituted by 1-2 adjacent connecting sites. The innervating layers taken into account are L2/3, L4, L5a, L5b and L6. The input field width corresponds to the distance between the most lateral and most medial connecting sites. The amplitude of EPSC at unitary connections is determined on monophasic currents. Of the 50 SPN included in the analysis of connectivity patterns in the barrel field, 35 were recorded for Amroune et al., 2025, using the same preparation and set-up.

#### Statistical analysis

We used non-parametric Wilcoxon, Mann-Whitney or Kruskal-Wallis tests, followed by Dunn-Sidak post hoc tests.

Differences between PV and SOM brains in starter and presynaptic cells centroid position investigated with anatomical tracing were assessed by permutation testing of the euclidean distance between group centroids (10000 permutations). To compare the relationship between cell count (centered on mean) and volume or mean maximal intercell distance in PVi and SOMi populations, a linear regression model including these variables (log_10_), interneuron type and their interaction was fitted to the data. The interaction term tested whether PVi and SOMi differed in these relationships.

Multivariate differences in connectivity patterns investigated with LSPS across neuronal types were evaluated using permutational multivariate analysis of variance (permanova - 999 permutations). The 5 parameters were standardized (z-score) prior to analysis. Data was downsampled to match the size of the smallest group. For determining the contribution of individual variables to group separation, each variable was permuted independently and the resulting reduction in the permanova pseudo-F statistic was quantified. For the distribution of functional connectivity across layers, row-wise differences between cell types were assessed using a two-sided cell-level permutation test with Benjamini Hochberg correction.

## Acknowledgments

This work was supported by grants from Agence Nationale de la Recherche: ANR-20-CE16-0002 and from the French government under the “France 2030” program via A*Midex (Initiative d’Excellence d’Aix-Marseille Université, AMX-19-IET-004, AMX-22-RE-AB-007/Neuradventure). The authors would like to thank Ede Rancz and Jyotika Bahuguna for careful reading of the manuscript and constructive feedback. We would like to thank interns who collaborated on the project (Lucie Limongi, Lucas Olivas, Michael Guillien and Abdel Ghazi). Helper and rabies viruses (AAV2.8.EEF1a.flex.GTc and AAV2.8.syn.flex.H2BG.N2cG, and RABV41) were a generous gift from Molly Strom, Viral VectorCore, The Francis Crick Institute, London, UK. We also thank all members of the team for support and discussions. Thanks to INMED core facilities, PBMC, animal facility and InMagic platform for help and support.

---

**Supplementary figure 1.**
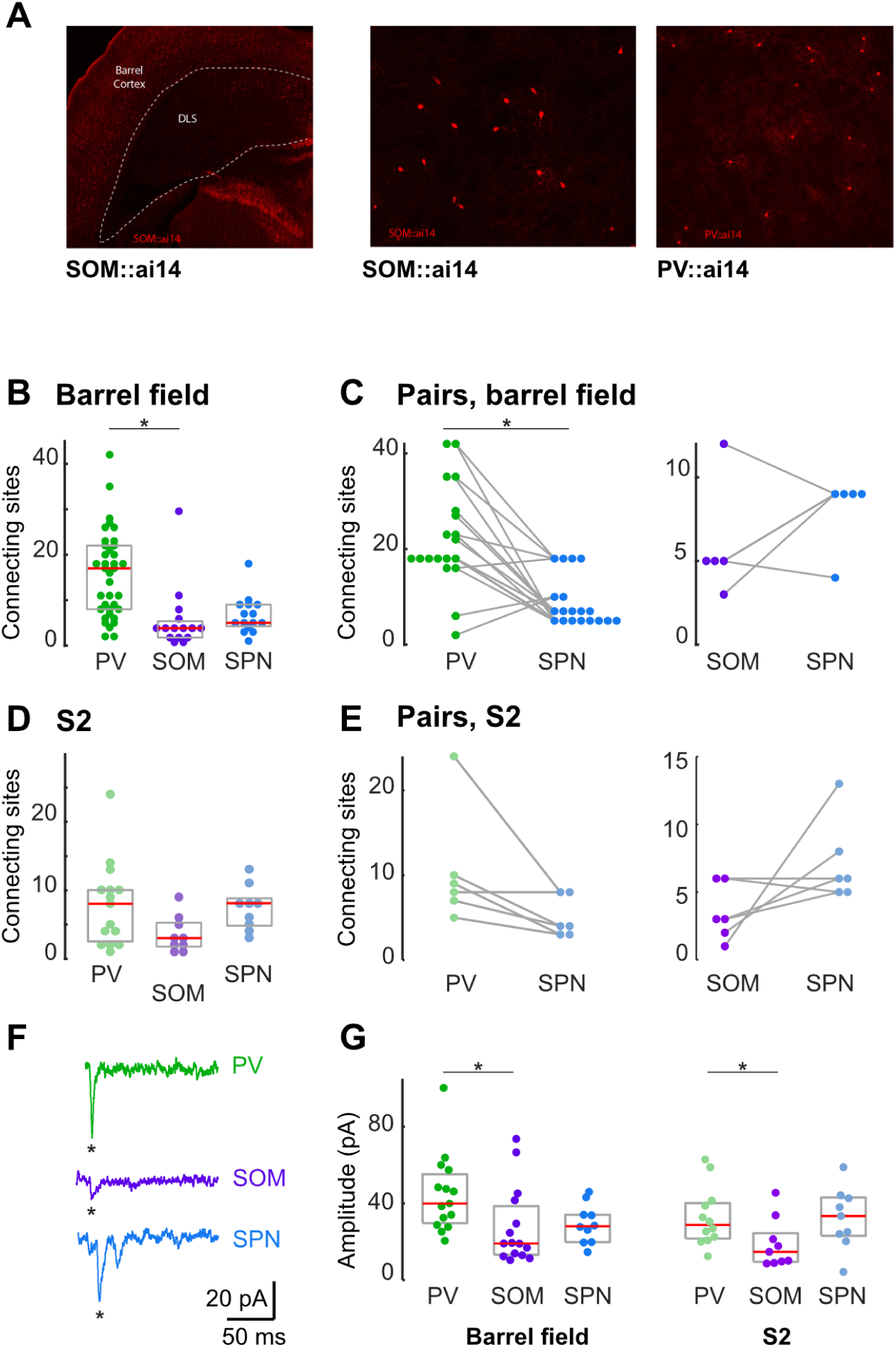
**A**: Labeled neurons in the DLS of SOM-cre::ai14 and PV-cre::ai14 mice at two magnifications. **B**: The number of connecting sites in PVi and SOMi LSPS connectivity maps (median and 25-75th percentiles). **C**, Paired comparison of the number of connecting cortical sites in PVi-SPN pairs (n = 19) and SOMi-SPN pairs (n = 5) recorded in the same slice. *, PV-SPN, p < 0.001, SOM-SPN, p = 0.25, Wilcoxon). **D** and **E**, same as in B and C. **F**, Example traces with unitary EPSC. *, analyzed events. **G**, Amplitude of evoked S1 evoked EPSCs from unitary connections recorded in PVi (n=31), SOMi (n=15) and SPN (n=10) and of S2 unitary EPSC recorded in PVi (n=12), SOMi (n=9) and SPN (n=9). Each symbol corresponds to a cell and may be the average of several EPSC. *, S1: p<0.005, PV-SOM p = 0.0163; S2: p <0.05, PV-SOM p = 0.0027; Kruskal-Wallis and Dunn-Sidak post hoc test.

